# No functional contribution of the gustatory receptor, Gr64b, co-expressed in olfactory sensory neurons of *Drosophila melanogaster*

**DOI:** 10.1101/2022.06.29.498097

**Authors:** Venkatesh Pal Mahadevan, Sofía Lavista-Llanos, Markus Knaden, Bill S. Hansson

**Affiliations:** Department of Evolutionary Neuroethology, Max Planck Institute for Chemical Ecology, Jena, Germany; CIFASIS-CONICET Franco-Argentine International Center for Information and Systems Sciences—National Council for Scientific and Technical Research, Rosario, Argentina

## Abstract

Chemosensation is essential for the survival of insects. Activities like searching for food, mating, and oviposition in the fruit fly, *Drosophila melanogaster* are to a great extent governed by chemical cues detected via olfaction and gustation. This chemical information is conveyed to higher brain centres via populations of diverse olfactory sensory neurons (OSNs) and gustatory sensory neurons (GSNs) expressing olfactory receptors (ORs) and gustatory receptors (GRs), respectively. ORs are exclusively expressed in the antenna and in the maxillary palps, while GRs are widely expressed in the labellum, tarsi, genitalia etc. Interestingly, 14 GRs were previously reported to be expressed in the antenna of *D. melanogaster*. However, the spatial expression pattern for all GRs and their functional role are still unclear. Recent data challenge the dogma that single OSNs express a single OR. In the present study, we studied the expression of 12 previously reported GRs among sensory structures on the fly antenna using the Gal4-UAS binary expression system. We observed antennal expression of nine out of the 12 reported. Out of these nine, consistent expression was only apparent for Gr64b, and we reconfirmed its presence in OSNs innervating three glomeruli in the antennal lobe. These glomeruli are known to be innervated by ab5A, ab5B and ab8A OSNs, respectively. Next, we generated double labelling crosses with Gr64b and observed co-expression of Gr64b with Or47a, which is expressed in the ab5B neuron. To elucidate the functional role of Gr64b co-expressed with Or47b, we challenged Or47a-expressing OSNs in wild type and Gr64b^-/-^ mutant flies with odor stimulation using the single sensillum recording technique in two satiation states (fed and starved). Notably, we did not observe any significant odor sensitivity or specificity changes in Gr64b mutants as compared to wild type flies. Taken together, our results reveal co-expression of GRs with ORs in olfactory sensory neurons, while the functional contribution of the GR in this context remains obscure.

## Introduction

Perception of multiple chemical cues is essential for the survival of insects. The fruit fly, *Drosophila melanogaster*, utilises such cues to find e.g., food sources (Stensmyr et al., 2003), suitable mates (Dweck et al., 2015) and oviposition substrates (Dweck et al., 2013) to name a few. The perception of chemical cues is carried out with the help of receptors expressed in peripheral sensory organs of the fly. In *D. melanogaster*, three gene families code for chemosensory receptors: olfactory receptors (ORs) (Clyne et al., 1999; Vosshall et al., 1999), gustatory receptors (GRs) (Montell, 2009)and ionotropic receptors (IRs) (Benton et al., 2009). Canonically, ORs are expressed in the antenna and maxillary palps, while GRs are expressed in multiple organs such as labellum, tarsi, wings, reproductive organs etc. (Depetris-Chauvin et al., 2015). Typically, ORs form a complex with the ubiquitous olfactory receptor co-receptor (Orco) and are responsible for the detection of a unique panel of odorants (Hallem and Carlson, 2006). On the other hand, GRs form heteromeric complexes and their ligand specificity changes depending upon the combination of GRs co-expressed (Jiao et al., 2008). Multiple studies have identified GRs that are required for tasting sweet and bitter compounds and describe diverse functional roles, such as inhibition of feeding and oviposition, inhibition of male to male courtship, etc. (Jiao et al., 2008; Moon et al., 2009; Watanabe et al., 2011; Lee et al., 2012; Freeman and Dahanukar, 2015; Fujii et al., 2015; Dweck and Carlson, 2020; Vernier et al., 2022). The first hint about non-canonical expression of GRs in the antenna of *D. melanogaster* came in the late 2000s, when two independent studies demonstrated that the perception of CO_2_ depends on Gr21a and Gr63a co-expressed in the ab1C neuron in the *D. melanogaster* antenna (Jones et al., 2007; Kwon et al., 2007). Follow-up studies using RNA-seq analysis and binary expression systems demonstrated expression of up to 14 GRs in the antenna, including GRs previously known to be involved in sugar and bitter taste perception (Thorne and Amrein, 2008; Ni et al., 2013; Menuz et al., 2014; Fujii et al., 2015). Based on these results, it was hypothesized that GRs might be co-expressed together with ORs, forming a heteromer and function in detection of novel ligands different from the native tuning profile of the corresponding OR (Fujii et al., 2015). However, apart from the deorphanization of Gr28bD expressed in the arista as being involved in avoidance of rapid increases in temperature (Ni et al., 2013), scant information is available regarding the functional significance of GR-OR co-occurrence in the same OSN. In other combinations, it has been reported that detection of sour taste via tarsal gustatory sensory neurons (GSNs) was mediated via co-expressed IRs (Chen and Amrein, 2017). Two recent studies also demonstrated overlapping expression of multiple gene families, with emphasis on IRs and ORs in the *D. melanogaster* antenna (Task et al., 2022) and in the yellow fever mosquito, *Aedes aegypti* (Younger et al., 2022). These studies are particularly important in demonstrating a functional role of IRs and ORs co-expressed in the OSNs. There are at least two possible explanations for such an overlap. First, such co-expression of multiple receptors might lead to heteromeric receptors that could ultimately accommodate novel ligands beyond the usual spectrum of each independent receptor. Secondly, such overlap could help in making the system redundant. If, by any natural incident, one receptor or receptor family fails then the other receptor could still maintain a partial functioning of the neurons. However, both of the studies mentioned above focused on the overlap of ORs and IRs with little focus on the involvement of the GR family.

In the present study, we focused on the co-expression of ORs and GRs in OSNs in the antenna of *D. melanogaster*. To generate a spatial expression map, we attempted to establish expression patterns of 12 GRs previously shown to be expressed in the antenna. Next we focused our attention on one specific sugar-detecting GR, Gr64b, due to its general expression pattern in the antenna and due to its co-expression with the well-characterised Or47a (also co-expressing Or33b) in the ab5B OSN. Using single sensillum recording technique, we investigated possible modulatory effects on odor sensitivity and specificity of the ab5B OSN in the absence of Gr64b. However, we did not observe any significant changes neither in odor sensitivity nor in specificity.

## Materials and methods

### Fly lines

Flies were reared at 25°C on standard cornmeal agar medium under 12h light: 12h dark photoperiod cycle. A mix of 7-15 day old males and females were used for immunohistochemistry and single sensillum recording (SSR) experiments. For starvation experiments, flies were starved for 20 hours with access to water *ad libitum* before being used for electrophysiology. Transgenic fly lines were obtained from the Bloomington fly stock centre with identities as listed in table 1.

**Table 1:** A list of all fly lines used during the experiments

| Fly line identity | Genotype | Bloomington ID |
| --- | --- | --- |
| Gr21a | Gr21a-gal4 | Originally from Vosshall lab |
| Gr22e | Gr22e-gal4 | BL57608 |
| Gr28bC | Gr28bC-gal4 | BL57618 |
| Gr28bD | Gr28bD-gal4 | BL57620 |
| Gr43a | Gr43a-gal4 | BL57636 |
| Gr61a | Gr61a-gal4 | Originally from Baker lab<br>(ID1190807) |
| Gr64a | Gr64a-gal4 | BL 57661 |
| Gr64b | Gr64b.mGFP (direct fusion<br>protein) | BL52630 |
| Gr64c | Gr64c-gal4 | BL57663 |
| Gr64f | Gr64f-gal4 | Originally from Baker lab<br>(ID1191452) |
| Gr93a | Gr93a-gal4 | BL57679 |
| mGFP | UAS-mGFP | BL5137 |
| mRFP | UAS-mRFP | BL27398 |
| Octuple GR mutant | $\Delta$ Gr5a, Gr64a-f, Gr61a null<br>mutant | Originally from Amrein lab |
| Or82a | Or82a-gal4 | BL 23126 |
| Or47a | Or47a-gal4 | BL 9982 |

### Chemical stimuli

All chemicals were purchased from Sigma Aldrich (Steinheim, Germany) with the highest purity (>98%). Geranyl acetate (CAS: 105-87-3) and pentyl acetate (CAS: 628-63-7) were used as diagnostic odors for ab5A and ab5B neurons, respectively. Both odors were serially diluted in hexane. A panel of 32 odors of ecological significance and representing diverse chemical groups were used to determine any changes in the odor tuning pattern of ab5A and B OSNs. A list of these 32 odors is available in table 2.

**Table 2:** List of chemicals used for identifying odor-tuning properties of neurons innervating ab5 sensillum class.

|  | Common name | CAS |
| --- | --- | --- |
| 1 | Hexane |  |
| 2 | Ethyl acetate | 141-78-6 |
| 3 | Ethyl lactate | 97-64-3 |
| 4 | CO<sub>2</sub> | Aspiration |
| 5 | Methyl salicylate | 119-36-8 |
| 6 | Methyl acetate | 79-20-9 |
| 7 | Ethyl-3-hydroxybutyrate | 5405-41-4 |
| 8 | ethyl hexanoate | 123-66-0 |
| 9 | 2-heptanone | 110-43-0 |
| 10 | geranyl acetate | 105-87-3 |
| 11 | pentyl acetate | 628-63-7 |
| 12 | 1-octen-3-ol | 3391-86-4 |
| 13 | ethyl benzoate | 93-89-0 |
| 14 | Ethyl crotonate | 623-70-1 |
| 15 | acetoin | 513-86-0 |
| 16 | 2 phenylalcohol | 60-12-8 |
| 17 | benzyl butyrate | 103-37-7 |
| 18 | 2-butanone | 78-93-3 |
| 19 | ethyl butanoate | 105-54-4 |
| 20 | isopropyl benzoate | 939-48-0 |
| 21 | methyl benzoate | 93-58-3 |
| 22 | 6-methyl-5-helpten-2-one | 110-93-0 |
| 23 | Hexyl acetate | 142-92-7 |
| 24 | Isopentyl propionate | 105-68-0 |
| 25 | 2-nonanol | 628-99-9 |
| 26 | 2-nonanone | 821-55-6 |
| 27 | Isopentyl acetate | 123-92-2 |
| 28 | 4-methylphenol | 106-44-5 |
| 29 | Acetophenone | 98-86-2 |
| 30 | methyl hexanoate | 106-70-7 |
| 31 | propyl acetate | 109-60-4 |
| 32 | nonanal | 124-19-6 |

### Electrophysiology: Single Sensillum Recordings

Single sensillum recordings were performed in order to establish the response profile of OSNs present in individual sensilla on the fly antenna. A 7-15-day old fly was immobilized in a 200 μl pipette tip and fixed on a glass side with laboratory wax. The funiculus (third antennal segment) was fixed in such a position that either the medial or posterior side faced the observer. Extracellular recordings were performed using electrochemically (3M KOH) sharpened tungsten electrodes by inserting a ground electrode in the eye and a recording electrode into the base of a sensillum using micromanipulators (Luigs and Nuemann SM-10). Sensilla were visualized with 1000x magnification using a binocular microscope (Olympus BX51W1). Signals were amplified (Syntech Uni-versal AC/DC Probe;www.syntech.nl), sampled (96000/s) and filtered (3kHz High-300Hz low, 50/60 Hz suppression) using a USB-IDAC. Neuronal activity was recorded using AutoSpike software (v3.7) for 3 sec pre and 10 sec post stimulus. The odor stimulus was delivered for 500 ms using a Pasteur pipette and was added to filtered and re-humidified air being constantly delivered to the fly antenna at 0.6 L/min through an 8mm id stainless steel tube ending 5mm from the antenna. Responses were established by calculating the change in spike frequency (spikes/s) 1 sec pre and post onset of the stimulus. Stimulus cartridges were prepared by pipetting 10 μl of the desired compound dissolved in hexane onto a filter paper (Rotilabo-round filters, type 601A, Carl Roth GmbH, Germany) with a diameter of 10 mm, which was then placed into a Pasteur pipette. Not more than two sensilla were recorded from each fly and odor stimuli were used for a maximum for two times. An inter-stimulus interval of at least 45 seconds was maintained and stimulation series always started with the lowest concentration and were then applied in an increasing order of concentrations.

### Immunohistochemistry

For whole-mount immunostaining, 7-15 day flies of mixed sexes were anesthetized on ice, and their antennae were dissected into cold *Drosophila* ringer solution with 0.1% Triton-X. The antennal mix consisted of specimens with only the third segment had been excised and those where both the second and third segments remaining. Dissected antennae (hereafter named “mix”) were fixed in 4% paraformaldehyde (with 0.1% Triton-X) for 1 hour on ice in a shaker. The mix was washed at least three times with 500 μl PT (1x PBS + 50 μl Triton) for 15 minutes each at room temperature. 500 μl Blocking solution (PT + 5% NGS) was added to the mix and incubated for 1 hour. Primary antibodies were diluted in 500 μl of fresh blocking solution and added to the mix followed by 48 hour incubation at 4°C. After the incubation, the mix was washed at least 3 times with PT, followed by addition of 500 μl blocking solution with 1-hour incubation. Secondary antibodies were diluted in 500 μl of fresh blocking solution and added to the mix followed by 48-hour incubation at 4°C. Lastly, the mix was washed at least 3 times with PT and mounted in Vectashield (Vector, Burlingame, CA, USA). Primary antibodies used were: mouse-anti-GFP (1:500), rabbit-anti-RFP (1:100) and rabbit-anti-orco (1:500), while secondary antibodies were: goat-anti-mouse (1:250) and goat-anti-rabbit (1:250). All antibodies used were purchased from Invitrogen (Invitrogen, Carlsbad, CA, USA).

### Statistical analysis

SSR traces were analysed using AutoSpike32 software 3.7 version (Syntech, NL 1998). Changes in action potential (spike count) were calculated by subtracting the number of spikes one sec before (spontaneous activity) from those elicited during one second after the onset of the stimulus. Spike changes for each concentration between wild type and GR mutant fly line were compared using unpaired, non-parametric t-tests with Mann Whitney test. Graphs were generated and statistical tests were performed using GraphPad Prism 9.1.1. Figures were constructed with Adobe Illustrator CS5 and Adobe Photoshop (Adobe system Inc.).

## Results

### Spatial expression map of Gustatory Receptors (GRs) in the antenna

Earlier studies have demonstrated expression of gustatory receptors (GRs) in the antenna of *D. melanogaster*. Here, we wanted to establish the spatial expression map of all previously reported GRs in the antenna. We took advantage of the binary expression system in *D. melanogaster*, combined 12 GR specific Gal4 drivers independently with membrane expressing GFP (UAS-mGFP) and checked for the spatial expression pattern of each of the GRs in the antenna (Figure 1A). We observed expression of two GRs (Gr93a and Gr28bC) in the second segment (Figure 1 H & J), five (Gr21a, Gr64c, Gr64b, Gr64f and Gr43a) in the third segment (Figure1 B, C, D, E and G), Gr28bD in the arista (Figure 1I) and Gr64e in both the second and third segments (Figure 1F). Using this data, we were able to generate a spatial expression map of these nine GRs in the antenna (Figure 1K). We proceeded to count the number of cells labelled by each GR driver (Figure 1L). Although several GRs were expressed in the antenna, only Gr64b displayed a consistent expression pattern throughout multiple replicates with a mean number of 20 GR64b positive cells (figure 1L). Therefore, we decided to focus our physiological investigations on Gr64b and proceeded to elucidate its role in olfaction during the next set of experiments.

**Figure 1:**
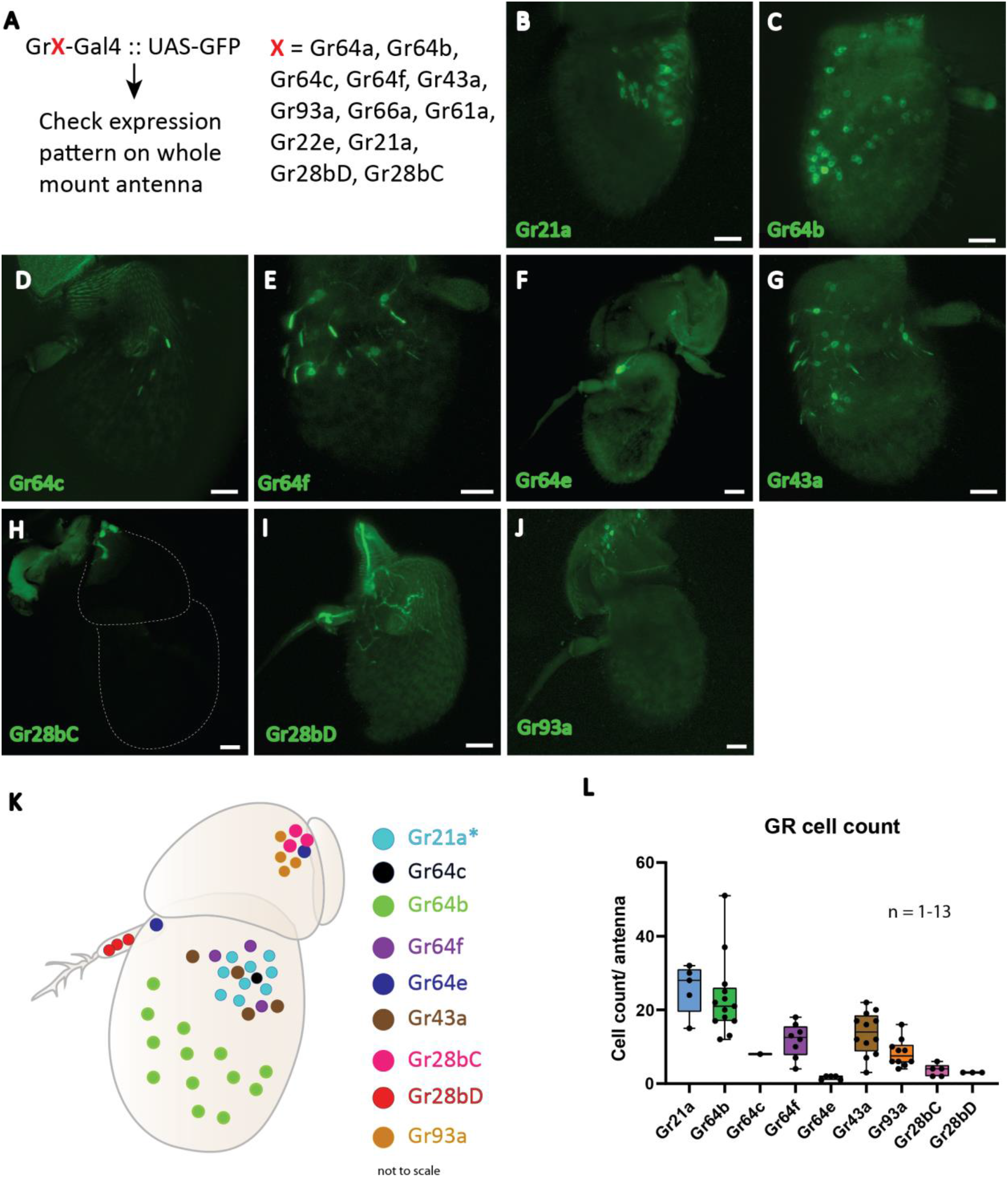
Spatial expression map of gustatory receptors expressed on *D. melanogaster* antenna. A: Schematic diagram of methodology where the Gal4-UAS binary expression system was used to check expression of 12 GRs. B-J: Expression patterns of individual gustatory receptors on the antenna with GFP. Scale bars = 20 µm K: Schematic representation of the spatial expression pattern of nine GRs on the antenna. Asterisk (*) denotes that the regions expressing Gr21a also co-express Gr63a in the same neuron and Gr10a in the neighboring neuron housed in the same sensilla. L: Cell counts of individual GR-Gal4 positive cells across multiple replicates observed positive for GFP signal.

### Gr64b is co-expressed with Or47a in ab5B neurons

To establish if Gr64b contributes to the olfactory coding in the cells where it is expressed, we first identified which physiological type of neurons express Gr64b. It is well known that Gr21a along with Gr63a, responsible for detecting CO_2_, is expressed in the ab1C neuron and that this neuron does not express the olfactory co-receptor Orco (Jones et al., 2007). We used this information as an internal positive control to verify the function of antibodies used (Supplementary fig 1). Next, we checked the co-expression pattern of Gr64b along with Orco by using a direct GFP fusion line (Gr64b.mGFP, Table 1) and an anti-Orco antibody. Although we observed co-expression of Gr64b in a few Orco positive cells, a significant population of Gr64b positive cells were Orco negative (Figure 2A). However, the cells positive for both Orco and Gr64b point out towards expression of Gr64b in OSNs. Next, we checked for the innervation pattern of axons of Gr64b-expressing OSNs in the antennal lobe. We could reconfirm that Gr64b positive neurons innervate VA6, DM3 and VM2 glomeruli in the antennal lobe along with canonical innervation in the suboesophageal ganglion as previously reported by Fujii et al., 2015 (Figure 2B). These three glomeruli, VA6, DM3 and VM3 have previously been shown to be innervated by OSNs expressing Or82a, Or47a (co-expressed with Or33b) and Or43b respectively. Therefore, we hypothesized that the small subpopulation of Gr64b and Orco co-expressing OSNs should also express at least one of the above-mentioned olfactory receptors. To start testing this hypothesis, we generated double cross fly lines with genotypes Or82a-Gal4 :: UAS-mRFP and Gr64b.mGFP. We did not observe any colocalization of Gr64b and Or82a (Figure 2C) but Gr64b positive neurons were located in the immediate vicinity of Or82a positive OSNs with adjacent dendritic projections (Figure 2D). The sensillum housing OSNs expressing Or82a and Or47a is named ab5. Next, we checked for the double cross with a genotype Or47a-Gal4:: UAS-mRFP and Gr64b.mGFP and could indeed observe colocalization (Figure E). Notably, only a small subpopulation of Gr64b-expressing cells also express Or47a (Figure E arrows) as many neuronal somata expressed only Gr64b (Figure E hand points). Taken together, we show that all Or47a-expressing neurons co-express Gr64b but the converse is not true (Figure E and F).

**Figure 2:**
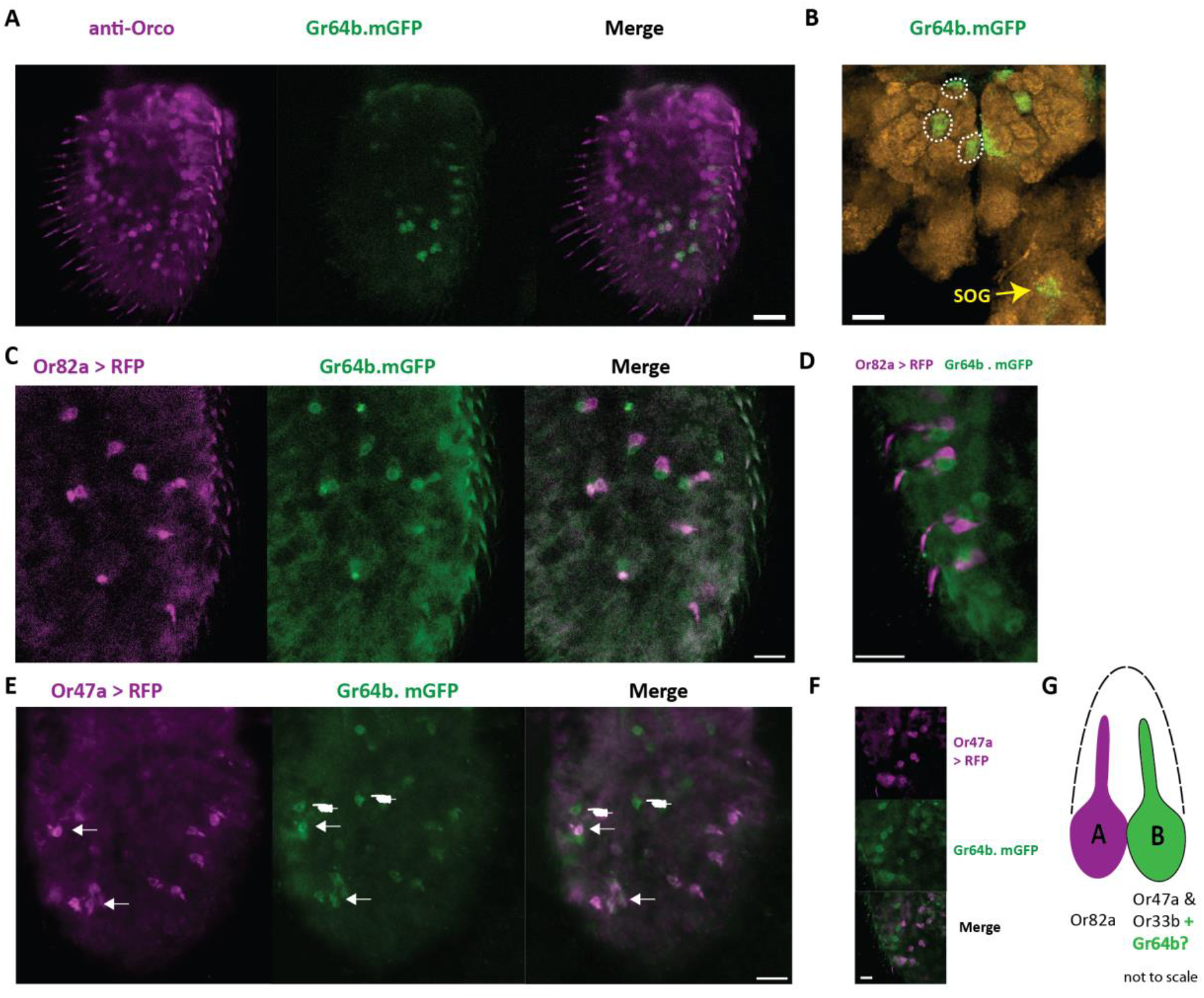
Gr64b is co-expressed with Or47a-Or33b in ab5B neuron in *D*.*melanogaster*. **A**: Staining of OSNs with anti-Orco antibody (magenta) and Gr64b expressing neurons tagged with GFP (green). Only a few cells are observed to be both Gr64b and Orco positive. Scale bar = 20 µm B: Innervation of three glomeruli (VA6, DM3 and VM2) by Gr64b positive neurons. Green color denotes Gr64b positive glomeruli while orange color denotes neuropils with nc82 staining. Innervation of the suboesophagal ganglion (SOG) denoted in yellow arrow. **C-D**: Double labelling crosses with Or82a-Gal4 driving expression of mRFP (magenta) and Gr64b tagged with GFP (green). No-colocalisation is observed. However, Or82a and Gr64b positive neurons appeared to be in the close vicinity of each other suggesting innervation of the same sensillum. Scale bars = 10 µm **E-F**: Double labelling crosses with Or47a-Gal4 driving expression of mRFP (magenta) and Gr64b tagged with GFP (green). Co-expression of Or47a was observed with Gr64b. All Or47a positive cells were also GFP positive (white arrow heads). However, a sub population with only Gr64b positive cells was also observed (white hand points). Scale bars = 10 µm G: A schematic model for the new non-canonical innervation pattern of the ab5 sensillum class in *D*.*melanogaster*.

### Functional implications of Gr64b on olfactory coding in ab5B olfactory sensory neurons

In the previous experiment, we showed that Gr64b often is co-expressed in Or47a expressing neurons. In *D. melanogaster*, Or47a expression corresponds to the B neuron innervating the ab5 sensillum class. We used single sensillum recording (SSR) technique to investigate if Gr64b contributes functionally to the odor sensitivity threshold and odor specificity of the Or47a-expressing ab5B OSN. We used a GR mutant line (referred to as sugar blind, (Yavuz et al., 2014) that carries a deletion of eight GRs, including Gr64b, all of which are involved in detection of sugars when expressed on the labellum. Even though Gr64b is not co-expressed with Or82a we also checked changes in the ab5A neuron, as a negative control. First, we tested if any changes in the odor sensitivity threshold of ab5B OSNs in the absence of Gr64b. We tested the SSR responses in two physiological states (fed and starved) so as to check for any feeding state dependent olfactory contribution of Gr64b in odor sensitivity. We measured the change in the number of action potentials (spikes) after odor stimulation, and we established the area under the spike frequency curve (figure 3A). The first analysis gives a measure of quantitative changes, while the latter reveals temporal changes in the firing dynamics of the neurons.

**Figure 3:**
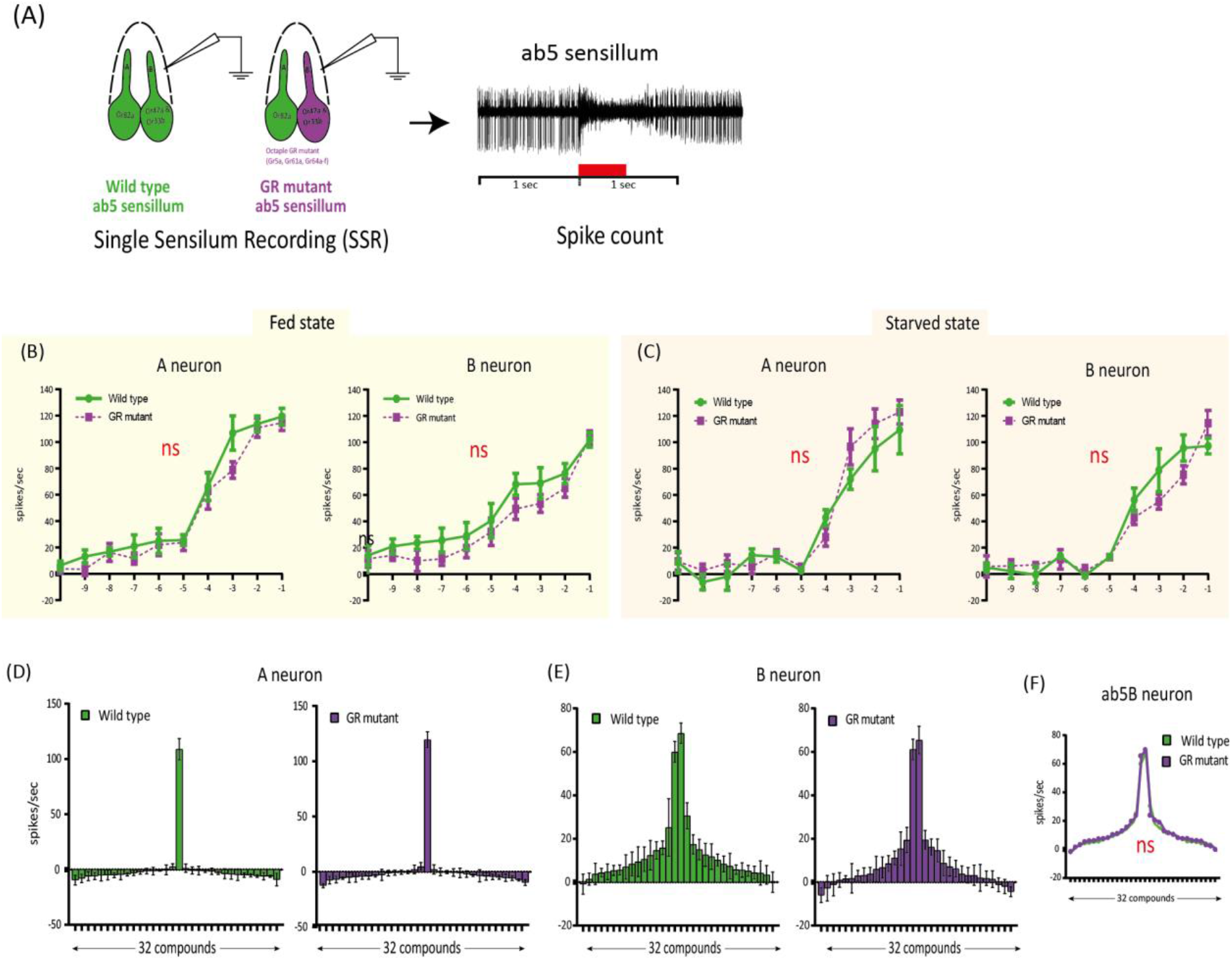
Single sensillum recording (SSR) experiments confirming no functional involvement of Gr64b in modulating odor sensitivity of ab5B neuron in *D*.*melanogaster*. **A**: Schematic representation of the methodology used to assess the functional role of Gr64b. SSR recordings were performed from the ab5 sensillum class in wild type CS fly and compared with an octuple GR mutant line (sugar blind fly, (Yavuz et al., 2014)) in two satiation states as fed and starved. Geranyl acetate and pentyl acetate were used as the best ligands for ab5A and ab5B neurons respectively. **B-C**: Dose response curves for both ab5A and ab5B neurons in fed and starved states with the green lines representing wild type CS fly and magenta line representing the octuple GR mutant fly. n = 4-6/ concentration. Unpaired, non-parametric t-tests with Mann Whitney test were performed at each concentration between two genotypes. No significant differences at each concentration were observed across experiments (p > 0.05, unpaired t-test). The sensitivity was not changed between two genotypes tested. **D-E**: F-H: Odor tuning sensitivity of both ab5A and ab5B neurons were checked with a panel of 32 odors representing diverse chemical classes in wild type and octuple GR mutant fly lines in fed state in case of ab5A and ab5B neurons respectively (Fig F-G) No significant shift in odor tuning curve in case of ab5B neuron was observed in absence of Gr64b (Fig H), n = 5.

The experiments revealed no significant differences in the odor sensitivity threshold between wild type flies and the octuple GR mutant line (Figure 3B-C). Both ab5A and ab5B neurons were equally sensitive to their best ligands (geranyl acetate and pentyl acetate, respectively) at low concentrations (10^−10^ v/v) in both fed and starved states. Nor were any significant differences in spike counts observed at any concentration tested (p > 0.05, unpaired, non-parametric t-tests with Mann Whitney test, Figure 3B-C).

Next, we studied changes in response specificity in ab5B OSNs lacking Gr64b. Only fed flies were used for these experiments. Once again, we did not observe any significant changes or shifts in odor tuning to a panel of 30 diverse ligands representing ecologically relevant odors of diverse chemical classes (Figure 3D-F, table 1). Taken together, our results suggest that Gr64b plays no significant role in modulating odor sensitivity or specificity in Or47a-expressing OSNs even though they are co-expressed in the very same neurons.

## Discussion

Canonical models of odorant receptor (OR) expression in *Drosophila melanogaster* have generally supported a one receptor type - one olfactory sensory neuron (OSN) class model. However, two recent studies challenge this model by showing expression of multiple chemoreceptor families in a single class of OSN in *D. melanogaster* (Task et al., 2022) and also in *Aedes aegypti* (Younger et al., 2022). Both of these studies demonstrated co-expression of the olfactory co-receptor (Orco) and multiple ionotropic co-receptors, Ir8a, Ir76b and Ir25a, in multiple OSN classes in the antenna and maxillary palps. However, the scope of these studies paid little attention to the expression and possible role of gustatory receptors (GRs) in the antenna. Although previous studies have reported expression of multiple GRs in the antenna of *D. melanogaster*, their functional role has not been elucidated (Thorne and Amrein, 2008; Fujii et al., 2015).

Here, we investigated if GRs contribute functionally to peripheral olfactory coding in *D. melanogaster*. We first tried to determine the expression patterns of the 12 gustatory receptors that have previously been reported to be expressed in the antenna of *D. melanogaster* (Thorne and Amrein, 2008; Menuz et al., 2014; Fujii et al., 2015) and generated a spatial expression map. Built on these results, we focused our functional study on Gr64b because of its consistent and broad expression pattern and its clear co-expression with Or47a, known to be expressed in ab5B neurons. Using single sensillum recording experiments in Gr64b mutant flies we did not observe any significant changes, neither in odor sensitivity, nor in odor turning properties of Or47a expressing neurons.

In our initial screening of expression patterns, we selected 12 GRs based on previous reports of expression of GRs in the antenna of *D. melanogaster* (Thorne and Amrein, 2008; Menuz et al., 2014; Fujii et al., 2015). Using the Gal4-UAS binary expression system we could only observe expression of nine out of the 12 GRs tested. Six of the nine GRs expressed can be roughly classified into two groups based on their canonical functions when expressed in the labellum. Gr64b, Gr64c, Gr64e, Gr64f and Gr43a in different heteromeric combinations have been reported to be involved in detection of sugars, glycerol and fatty acids (Jiao et al., 2008; Wisotsky et al., 2011; Fujii et al., 2015; Kim et al., 2018) while Gr93a is involved in detection of the bitter tastant caffeine when expressed in combination with other bitter GRs (Lee et al., 2009). Two GRs, Gr28bD and Gr21a, have been reported to be involved in rapid warmth sensing and CO_2_ perception respectively while Gr28bC still remains functionally orphan (Jones et al., 2007; Kwon et al., 2007; Ni et al., 2013).

We observed nine out of the twelve tested GRs. Labeling of neurons using binary expression systems greatly depend on the construction of the transgenic line (positional effect). It is thus possible that some essential regulatory elements for the expression could have been replaced during the genetic insertion (Yavuz et al., 2014; Task et al., 2022). This is specifically true in case of GRs from the GR64 cluster due to the complex arrangement of regulatory elements, sometimes also within the locus (Fujii et al., 2015). This limitation of the Gal4-UAS system was resolved by generation of LexA lines, and it could e.g. be shown that Gr64f has a broad expression in the antenna as well as in the maxillary palps of *D. melanogaster* (Fujii et al., 2015). Furthermore, a broad expression pattern for Gr64b as well as innervation of Gr64b expressing OSNs in the antennal lobe was reported. However, the authors did not investigate the functional role of any of these non-canonically expressed GRs (Fujii et al., 2015). In our study, using a direct fusion protein (Gr64b.mGFP), we observed a consistent and broad expression pattern for Gr64b. Interestingly, we observed expression of Gr64e only in a small subset of neurons in the funiculus in close vicinity to the base of the arista. This subset of neurons was Orco negative (supplementary fig 2) however, we cannot rule out the possibility that this subset of neurons could also express one or more IR coreceptor. When expressed in the proboscis, Gr64e has been reported to be involved in the perception of glycerol and fatty acids along with Gr64b (Wisotsky et al., 2011; Kim et al., 2018). We did not check if Gr64b and Gr64e were co-expressed. However, we did not observe a particular cluster of Gr64b positive cells at the base of the arista where Gr64e is expected. It is thus likely that these cells represent an independent set of neurons positive for Gr64e. The arista of *D. melanogaster* houses six cells out of which three are responsible for cold and rapid heat sensing. These cells express Ir25a (along with other IRs) and Gr28bD, respectively (Ni et al., 2013; Budelli et al., 2019). We observed efferents of Gr64e expressing neurons leading to the arista. A possibility open for future investigation is that Gr64e would be involved in modulating the function of Gr28bD. The second segment of the antenna (pedicle) houses the auditory center, Johnston’s organ, and has been demonstrated to be innervated by Gr28bC- and Gr68a-epressing neurons (Ejima and Griffith, 2008; Thorne and Amrein, 2008). Silencing of Gr68a-expressing neurons resulted in hampering the courtship behavior in *D. melanogaster* and thereby demonstrated another non-canonical expression and functional involvement of GRs when expressed in the auditory center of the antenna. We observed expression of Gr93a, Gr28bC and a few cells labelled with Gr64e on the second segment. Future investigations should establish if Gr28bC, Gr64e and Gr93a are generally expressed in the same set of neurons and if they contribute to the auditory and specifically mating behaviors of the fly.

Gr64b-expressing OSNs innervate three glomeruli in the antennal lobe (Fujii et al., 2015). Our results confirm this observation. We expected that due to the innervation of both the VA6 and the DM3 glomeruli, Gr64b should be co-expressed in both Or82a- and Or47a-expressing OSNs, known to innervate these two glomeruli (Couto et al., 2005). However, we observed co-expression of Gr64b only with Or47a (figure 2E), while cells positive for Gr64b and Or82a were separate but in close vicinity, suggesting innervation of the same sensillum (figure 2D). This mismatch raises a question regarding labelling of the VA6 glomerulus (innervated by Or82a) by Gr64b-expressing OSNs even though there is no co-expression at the OSN level. Such mismatch of receptor expression and glomerular innervation has been studied in detail in the recent two papers showing co-expression of Orco and IR25a (Task et al., 2022; Younger et al., 2022). In case of *D. melanogaster*, innervation of multiple glomeruli innervated by Ir25a positive neurons was observed (Task et al., 2022). It was earlier shown that the V glomerulus is innervated only by neurons expressing Gr21a and Gr63a, which are involved in detection of CO_2_ (Jones et al., 2007; Kwon et al., 2007). However, surprisingly, it was in the recent study by Task et al. (2022) observed that a few Orco and Ir25a positive neurons also innervate the V glomerulus sparsely, pointing to a wider possibility of overlap between these three gene families, but which could be revealed only after using more effective expression systems (Task et al., 2022). Similarly, the VM6 glomerulus, which was earlier considered a singular entity, was shown to be innervated by subpopulations of different neuron classes. A subdomain was innervated by a new type of OSNs expressing an Rh50 ammonium transporter protein (Vulpe et al., 2021) and also by Ir25a positive OSNs (Task et al., 2022). Further, it was shown that the VA6 glomerulus (corresponding to Or82a) was innervated by OSNs expressing multiple coreceptors, including Orco, Ir25a, Ir8a and Ir76b, while, interestingly, the DM3 (corresponding to Or47a and Or33b) was innervated only by Orco positive neurons. It is also interesting to note that Or47a-expressing neurons do not co-express any of these widely expressed IR co-receptors, yet express Gr64b. We did not check if there is co-expression of IR coreceptors and Gr64b. However, it would be interesting and informative for the future to check if these two gene families overlap in the antenna of *D. melanogaster*.

One question that is crucial to our understanding of the fly olfactory system is thus whether non-canonically co-expressed GRs contribute functionally to peripheral olfactory coding. Till date, a functional role has been assigned only to three antennal GRs (Gr21a, Gr63a and Gr28bD). Amongst those, Gr21a and Gr63a are co-expressed in the ab1C neuron in the funiculus and are involved in detection of CO_2_ (Jones et al., 2007; Kwon et al., 2007) while Gr28bD is expressed in the arista and has been shown to be involved in rapid heat sensing (Ni et al., 2013). Co-expression of Gr10a with Or10a in the ab1D neuron in the funiculus has also been reported without any further study regarding its role in odor perception (Jones et al., 2007). In the study by Task et al. (2022, see above) the authors checked for functional changes in OSNs co-expressing Ir25a using SSR technique. They found small but significant changes in a few sensillum classes in Ir25a mutant genotypes. However, functional changes were different for individual OSN types, as e.g. depression of spike numbers in case of ab3A OSN type when stimulated with 1-octen-3-ol, while increase in spike numbers in the pb1A and pb3A types when stimulated with the same compound. This demonstrates that removing Ir25a affects functionality in different OSN types independently. However, if this leads to any behavioral changes is yet to be explored. What ecological benefit could such overlap confer to an insect? A possible answer to this question lies in the formation of redundancy in the olfactory system and was demonstrated in another recent study in *Aedes aegypti* (Younger et al., 2022). There have been multiple attempts to generate genetically modified strains of anthropophilic mosquitoes by masking human odors. This was attempted either by deleting receptors involved in CO_2_ perception or by those involved in human odor sensing (Degennaro et al., 2013; McMeniman et al., 2014; Raji et al., 2019). However, deletion of either of these receptors did not completely hamper the host seeking abilities of mosquitoes under semi field conditions. A possible explanation to these observations was recently presented, revealing an extensive overlap between multiple gene families in OSNs of the *Aedes aegypti* antenna, maxillary palps and, to some extent, proboscis (Younger et al., 2022). The authors showed co-expression of Orco and Ir25a in multiple OSNs. The striking feature of these recent findings is that the olfactory system contains subsets of neurons positive for either only Orco or only Ir25a, while some neurons are positive for both Orco and IRs. This holds true also in the case of CO_2_ sensing Gr1-3-expressing neurons. Such redundancy is probably one of the key factors explaining the highly successful host seeking seen in mosquitoes, even in the absence of Orco. In this context, we were curious if there is any functional significance for the overlap between GR64b and Or47a expression and investigated if Gr64b has a role in odor sensitivity as well as in the tuning profile of Or47a-expressing OSNs on the *D. melanogaster* antenna. The odor tuning curve of an OSN is primarily determined by the ORs (Hallem and Carlson, 2006) but another study demonstrated amplification of Or67d neuronal responses in the combination of another receptor family protein, ppk25 (Ng et al., 2019). We did not observe any functional involvement of Gr64b in modulating odor sensitivity of Or47a OSNs in case of change in sensitivity threshold or changes in odor tuning specificity, suggesting no functional involvement at least in the odor detection characteristics of Or47a. We cannot yet rule out the possibility of other non-olfactory functional roles of Gr64b. Other non-canonical roles for Gr64b have also been suggested as one of the downstream factors for fatty acid sensing via Gr64e (Kim et al., 2018). Recently, a role of Gr64 cluster genes was demonstrated in proteostasis and is crucial for the survival of ribosomal protein mutant epithelial cells in *D. melanogaster* (Baumgartner et al., 2021). Lastly, a pleiotropic role of GRs has been well documented in case of expression in the fly brain, gastrointestinal tract and reproductive sites (Park and Kwon, 2011; Miyamoto and Amrein, 2013; Vernier et al., 2022). Therefore, it can be hypothesized that Gr64b may not perform a chemosensory role but rather act as a GPCR involved in cellular functions when expressed in ab5B neurons.

## Ethics Statement

This study on the fruit fly *Drosophila melanogaster* was performed in Germany where the research on invertebrates does not require a permit from a committee that approves animal research.

## Author Contributions

S-LL, MK and BH conceived the project. VPM and S-LL designed the experiments. VPM conducted the experiments, made the figures and data analysis. All authors discussed the results and wrote the article.

## Acknowledgments

We thank Regina Stieber, Silke Trautheim, Sin Prelic, Carolin Hoyer and Dr. Veit Grabe for comments, discussions and technical support.

## References

Baumgartner, M. E., Kucinski, I., and Piddini, E. (2021). Ribosome protein mutant cells rely on the GR64 cluster of gustatory receptors for survival and proteostasis in Drosophila. bioRxiv, 2021.05.10.443504. doi: 10.1101/2021.05.10.443504.

Benton, R., Vannice, K. S., Gomez-Diaz, C., and Vosshall, L. B. (2009). Variant Ionotropic Glutamate Receptors as Chemosensory Receptors in Drosophila. Cell 136, 149–162. doi: 10.1016/j.cell.2008.12.001.

Budelli, G., Ni, L., Berciu, C., van Giesen, L., Knecht, Z. A., Chang, E. C., et al. (2019). Ionotropic Receptors Specify the Morphogenesis of Phasic Sensors Controlling Rapid Thermal Preference in Drosophila. Neuron 101, 738–747.e3. doi: 10.1016/J.NEURON.2018.12.022.

Chen, Y., and Amrein, H. (2017). Ionotropic Receptors Mediate Drosophila Oviposition Preference through Sour Gustatory Receptor Neurons. Curr. Biol. 27, 2741–2750.e4. doi: 10.1016/J.CUB.2017.08.003.

Clyne, P. J., Warr, C. G., Freeman, M. R., Lessing, D., Kim, J., and Carlson, J. R. (1999). A novel family of divergent seven-transmembrane proteins: Candidate odorant receptors in Drosophila. Neuron 22, 327–338. doi: 10.1016/S0896-6273(00)81093-4.

Couto, A., Alenius, M., and Dickson, B. J. (2005). Molecular, Anatomical, and Functional Organization of the Drosophila Olfactory System. Curr. Biol. 15, 1535–1547. doi: 10.1016/j.cub.2005.07.034.

Degennaro, M., McBride, C. S., Seeholzer, L., Nakagawa, T., Dennis, E. J., Goldman, C., et al. (2013). orco mutant mosquitoes lose strong preference for humans and are not repelled by volatile DEET. Nat. 2013 4987455 498, 487–491. doi: 10.1038/nature12206.

Depetris-Chauvin, A., Galagovsky, D., and Grosjean, Y. (2015). Chemicals and chemoreceptors: Ecologically relevant signals driving behavior in Drosophila. Front. Ecol. Evol. 3, 41. doi: 10.3389/fevo.2015.00041.

Dweck, H. K., and Carlson, J. R. (2020). Molecular Logic and Evolution of Bitter Taste in Drosophila. Curr. Biol. 30, 17–29.e4. doi: 10.1016/j.cub.2019.11.005.

Dweck, H. K. M., Ebrahim, S. A. M., Kromann, S., Bown, D., Hillbur, Y., Sachse, S., et al. (2013). Olfactory Preference for Egg Laying on Citrus Substrates in Drosophila. Curr. Biol. 23, 2472–2480. doi: 10.1016/J.CUB.2013.10.047.

Dweck, H. K. M., Ebrahim, S. A. M., Thoma, M., Mohamed, A. A. M., Keesey, I. W., Trona, F., et al. (2015). Pheromones mediating copulation and attraction in Drosophila. Proc. Natl. Acad. Sci. U. S. A. 112, E2829–E2835. doi: 10.1073/PNAS.1504527112.

Ejima, A., and Griffith, L. C. (2008). Courtship Initiation Is Stimulated by Acoustic Signals in Drosophila melanogaster. PLoS One 3, e3246. doi: 10.1371/JOURNAL.PONE.0003246.

Freeman, E. G., and Dahanukar, A. (2015). Molecular neurobiology of Drosophila taste. Curr. Opin. Neurobiol. 34, 140–148. doi: 10.1016/j.conb.2015.06.001.

Fujii, S., Yavuz, A., Slone, J., Jagge, C., Song, X., and Amrein, H. (2015). Drosophila sugar receptors in sweet taste perception, olfaction, and internal nutrient sensing. Curr. Biol. 25, 621–627. doi: 10.1016/j.cub.2014.12.058.

Hallem, E. A., and Carlson, J. R. (2006). Coding of Odors by a Receptor Repertoire. Cell 125, 143–160. doi: 10.1016/J.CELL.2006.01.050/ATTACHMENT/4FFF1BF0-C0FF-40DB-AB19-97AA0CC4C01B/MMC1.PDF.

Jiao, Y., Moon, S. J., Wang, X., Ren, Q., and Montell, C. (2008). Gr64f Is Required in Combination with Other Gustatory Receptors for Sugar Detection in Drosophila. Curr. Biol. 18, 1797–1801. doi: 10.1016/j.cub.2008.10.009.

Jones, W. D., Cayirlioglu, P., Grunwald Kadow, I., and Vosshall, L. B. (2007). Two chemosensory receptors together mediate carbon dioxide detection in Drosophila. Nature 445, 86–90. doi: 10.1038/nature05466.

Kim, H., Kim, H., Kwon, J. Y., Seo, J. T., Shin, D. M., and Moon, S. J. (2018). Drosophila Gr64e mediates fatty acid sensing via the phospholipase C pathway. PLoS Genet. 14. doi: 10.1371/journal.pgen.1007229.

Kwon, J. Y., Dahanukar, A., Weiss, L. A., and Carlson, J. R. (2007). The molecular basis of CO2 reception in Drosophila. Proc. Natl. Acad. Sci. U. S. A. 104, 3574–3578. doi: 10.1073/pnas.0700079104.

Lee, Y., Kang, M. J., Shim, J., Cheong, C. U., Moon, S. J., and Montell, C. (2012). Gustatory receptors required for avoiding the insecticide L-canavanine. J. Neurosci. 32, 1429–1435. doi: 10.1523/JNEUROSCI.4630-11.2012.

Lee, Y., Moon, S. J., and Montell, C. (2009). Multiple gustatory receptors required for the caffeine response in Drosophila. Proc. Natl. Acad. Sci. U. S. A. 106, 4495–4500. doi: 10.1073/pnas.0811744106.

McMeniman, C. J., Corfas, R. A., Matthews, B. J., Ritchie, S. A., and Vosshall, L. B. (2014). Multimodal Integration of Carbon Dioxide and Other Sensory Cues Drives Mosquito Attraction to Humans. Cell 156, 1060–1071. doi: 10.1016/J.CELL.2013.12.044.

Menuz, K., Larter, N. K., Park, J., and Carlson, J. R. (2014). An RNA-Seq Screen of the Drosophila Antenna Identifies a Transporter Necessary for Ammonia Detection. PLoS Genet. 10. doi: 10.1371/journal.pgen.1004810.

Miyamoto, T., and Amrein, H. (2013). Diverse roles for the Drosophila fructose sensor Gr43a. http://dx.doi.org/10.4161/fly.27241 8, 19–25. doi: 10.4161/FLY.27241.

Montell, C. (2009). A taste of the Drosophila gustatory receptors. Curr. Opin. Neurobiol. 19, 345–353. doi: 10.1016/j.conb.2009.07.001.

Moon, S. J., Lee, Y., Jiao, Y., and Montell, C. (2009). A Drosophila Gustatory Receptor Essential for Aversive Taste and Inhibiting Male-to-Male Courtship. Curr. Biol. 19, 1623–1627. doi: 10.1016/J.CUB.2009.07.061/ATTACHMENT/3AD001B8-B776-4E68-BF54-A290EF736985/MMC1.PDF.

Ng, R., Salem, S. S., Wu, S. T., Wu, M., Lin, H. H., Shepherd, A. K., et al. (2019). Amplification of Drosophila Olfactory Responses by a DEG/ENaC Channel. Neuron 104, 947–959.e5. doi: 10.1016/J.NEURON.2019.08.041.

Ni, L., Bronk, P., Chang, E. C., Lowell, A. M., Flam, J. O., Panzano, V. C., et al. (2013). A gustatory receptor paralogue controls rapid warmth avoidance in Drosophila. Nature 500, 580–584. doi: 10.1038/nature12390.

Park, J. H., and Kwon, J. Y. (2011). Heterogeneous Expression of Drosophila Gustatory Receptors in Enteroendocrine Cells. PLoS One 6, e29022. doi: 10.1371/JOURNAL.PONE.0029022.

Raji, J. I., Melo, N., Castillo, J. S., Gonzalez, S., Saldana, V., Stensmyr, M. C., et al. (2019). Aedes aegypti Mosquitoes Detect Acidic Volatiles Found in Human Odor Using the IR8a Pathway. Curr. Biol. 29. doi: 10.1016/j.cub.2019.02.045.

Stensmyr, M. C., Giordano, E., Balloi, A., Angioy, A. M., and Hansson, B. S. (2003). Novel natural ligands for Drosophila olfactory receptor neurones. J. Exp. Biol. 206, 715–724. doi: 10.1242/JEB.00143.

Task, D., Lin, C.-C., Vulpe, A., Afify, A., Ballou, S., Brbic, M., et al. (2022). Chemoreceptor co-expression in Drosophila melanogaster olfactory neurons. Elife 11, 1–69. doi: 10.7554/eLife.72599.

Thorne, N., and Amrein, H. (2008). Atypical expression of Drosophila gustatory receptor genes in sensory and central neurons. J. Comp. Neurol. 506, 548–568. doi: 10.1002/cne.21547.

Vernier, C., Zelle, K. M., Leitner, N., Liang, X., Halloran, S., Millar, J. G., et al. (2022). A pleiotropic chemoreceptor facilitates the functional coupling of pheromone production and perception. bioRxiv, 2022.01.10.475668. doi: 10.1101/2022.01.10.475668.

Vosshall, L. B., Amrein, H., Morozov, P. S., Rzhetsky, A., and Axel, R. (1999). A Spatial Map of Olfactory Receptor Expression in the Drosophila Antenna. Cell 96, 725–736. doi: 10.1016/S0092-8674(00)80582-6.

Vulpe, A., Kim, H. S., Ballou, S., Wu, S. T., Grabe, V., Nava Gonzales, C., et al. (2021). An ammonium transporter is a non-canonical olfactory receptor for ammonia. Curr. Biol. 31, 3382–3390.e7. doi: 10.1016/J.CUB.2021.05.025.

Watanabe, K., Toba, G., Koganezawa, M., and Yamamoto, D. (2011). Gr39a, a Highly diversified gustatory receptor in drosophila, has a role in sexual behavior. Behav. Genet. 41, 746–753. doi: 10.1007/s10519-011-9461-6.

Wisotsky, Z., Medina, A., Freeman, E., and Dahanukar, A. (2011). Evolutionary differences in food preference rely on Gr64e, a receptor for glycerol. Nat. Neurosci. 14, 1534–1541. doi: 10.1038/NN.2944.

Yavuz, A., Jagge, C., Slone, J., and Amrein, H. (2014). A genetic tool kit for cellular and behavioral analyses of insect sugar receptors. Fly (Austin). 8, 189–196. doi: 10.1080/19336934.2015.1050569/SUPPL_FILE/KFLY_A_1050569_SM3008.PDF.

Younger, M. A., Herre, M., Goldman, O. V., Lu, T.-C., Caballero-Vidal, G., Qi, Y., et al. (2022). Non-Canonical Odor Coding in the Mosquito. bioRxiv, 2020.11.07.368720. doi: 10.1101/2020.11.07.368720.

