## Supplementary file1 for "No functional contribution of the gustatory receptor, Gr64b, co-expressed in olfactory sensory neurons of *Drosophila melanogaster*"

### Supplementary material

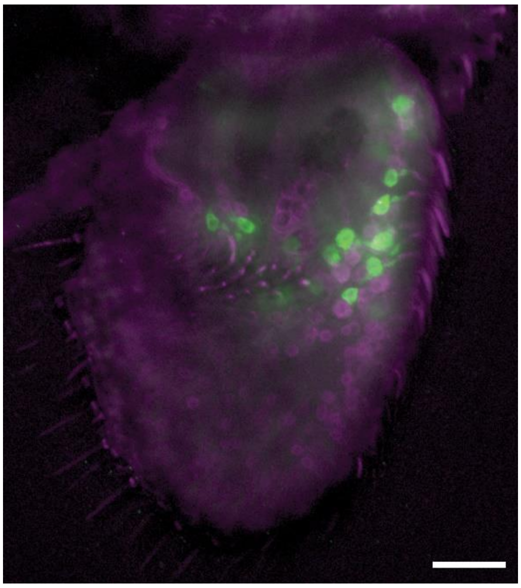

**Supplementary figure 1**: Double staining for Gr21a and Orco. Double labelling crosses with Gr21a-Gal4 driving expression of mGFP (green) and anti-Orco antibody (magenta) was used to check the working of anti-Orco antibody.

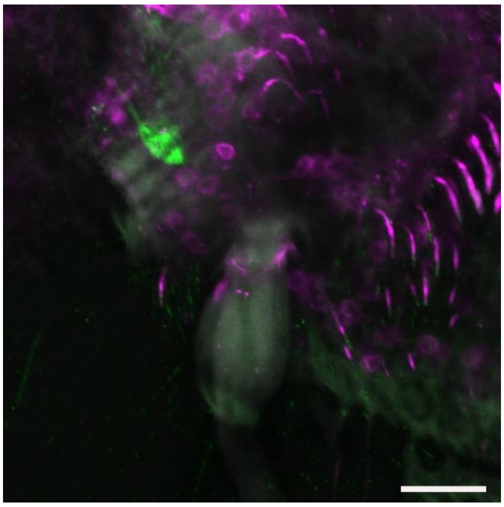

**Supplementary figure 2**: Double staining for Gr64e and Orco. Double labelling crosses with Gr64e-Gal4 driving expression of mGFP (green) and anti-Orco antibody (magenta) was used to check for co-expression of Gr64e in Orco positive cells.
